# Molecular Insights into Fungal Innate Immunity Using the *Neurospora crassa - Pseudomonas syringae* Model

**DOI:** 10.1101/2025.01.22.633611

**Authors:** Frances Grace Stark, Mari Torii-Karch, Sudyut Yuvaraj, Lucas Bonometti, Pierre Gladieux, N. Louise Glass, Ksenia Krasileva

**Affiliations:** University of California Berkeley, Department of Plant and Microbial Biology.; University of California Berkeley, Center for Computational Biology.; Plant Health Institute Montpellier, INRAE, CIRAD, Institut Agro, IRD, Université de Montpellier, France

## Abstract

Recent comparative genomics and mechanistic analyses support the existence of a fungal immune system. Fungi encode genes with features similar to non-self recognition systems in plants, animals, and bacteria. However, limited functional or mechanistic evidence exists for the surveillance-system recognition of heterologous microbes in fungi. We found that *Neurospora* species coexist with *Pseudomonas* in their natural environment. We leveraged two model organisms, *Neurospora crassa* and *Pseudomonas syringae* DC3000 (PSTDC3000) to observe immediate fungal responses to bacteria. PSTDC3000 preferentially surrounds *N. crassa* cells on a solid surface, causing environmental dependent growth responses, bacterial proliferation and varying fungal fitness. Specifically, the Type III secretion system (T3SS) ΔhrcC mutant of PSTDC3000 colonized *N. crassa* hyphae less well. To dissect initial cellular signaling events within the population of germinated asexual spores (germlings), we performed transcriptomics on *N. crassa* after PSTDC3000 inoculation. Upon contact with live bacteria, a subpopulation of fungal germlings initiate a response as early as ten minutes post-contact revealing transcriptional differentiation of Reactive Oxygen Species (ROS) mechanisms, trace metal warfare, cell wall remodeling dynamics, multidrug-efflux transporters, secondary metabolite synthesis, and excretion. We dissected mutants of plausible receptors, signaling pathways, and responses that *N. crassa* uses to detect and mount a defense against PSTDC3000 and found seven genes that influence resistant and susceptibility phenotypes of *N. crassa* to bacterial colonization. Mutants in genes encoding a ctr copper transporter (*tcu-1*), ferric reductase (*fer-1*), superoxide reductase (*sod-2*), multidrug resistance transporter (*mdr-6*), a secreted lysozyme-Glycoside hydrolase (*lyz*) and the Woronin body tether leashin (NCU02793, *lah-1* and *lah-2*) showed a significant reduction of growth in the presence of bacteria, allowing the bacteria to fully take over the fungal mycelium faster than wildtype. In this study we provide a bacterial-fungal model system within Dikarya that allows us to begin to dissect signaling pathways of the putative fungal immune system.

## Background

Research on fungal defenses and microbial interactions began in the twentieth century with many years of antimicrobial discovery, including Alexander Fleming’s famous penicillin discovery from a *Penicillium* and *Staphylococci* coculture [1]. Interactions between fungi and other microbes have been studied in isolation or in larger ecological contexts ever since, usually within microbial systems present in agriculture and important to human health [2–4]. Fungal-microbe literature has led to hypotheses in the microbial interaction field on how fungi generally defend themselves from microbial colonization, often citing nutrient starvation, trace-metal starvation [5], antimicrobial molecule synthesis [6,7], niche protection by priority effect, long distance diffusible signaling [8,9], RNA interference systems for viruses [10,11], and Repeat Induced Point mutation (RIP) protecting from transposons and other selfish genetic elements [12,13].

Bacteria and fungi compete to degrade plant matter. While fungi are generally considered to produce antimicrobials to protect themselves and their niche from competing bacteria, there is mounting evidence that antimicrobial synthesis is induced by the presence of bacteria, both internally and externally, and that these responses largely overlap, regardless of bacterial species [5,14–16]. Antimicrobial defenses play an important role in resistance mechanisms induced by specific surveillance-systems in plants, animals, and bacteria. In Fungi, this concept was recently discussed in Künzler et al. (2018) [6], describing fungal defense as either autonomous or antagonist-dependent. It is still unknown how these processes are induced, attuned, conserved, and involved in resistance/susceptibility phenotypes of fungi to bacteria. It is hypothesized that a sensor would start a signalling cascade leading to a specific, protective response. Yet limited evidence exists on receptors and signaling cascades dictating fungal responses to bacteria.

Within a comparative immunology framework [17], fungi should have conserved and specific ways of responding to bacterial pressure via specific receptors/sensors. This gap in knowledge exists because traditional phenotypes measured in immunology, such as known transcription factors, various types of cell death, immune system products/response, and fitness outcomes, are only starting to be compared to the cellular responses already uncovered in fungal-bacterial systems [17–20]. Fungal-bacterial study systems with well-defined effects of bacterial presence on fungal fitness, resistance or susceptibility coupled with characterized receptors and responses are needed to fill this gap [17,21]. Only one example exists of a fungal-bacterial interaction system with a defined receptor: *Candida albicans* recognizes *Pseudomonas aeruginosa* peptidoglycan with the Leucine Rich Repeat (LRR) domain of adenylyl cyclase Cyr1p, which is responsible for initiating *C. albicans* pseudohyphal growth, and mediates a symbiosis with a more virulent *P. aeruginosa* and *C. albicans* biofilm in human lungs [22].

Indirect evidence of receptors/sensors mediating fungal perception of bacteria are scattered throughout the literature. Some of the well documented cellular responses fungi have towards bacteria are: changing the profile of metabolism, transcription, translation, antimicrobial synthesis/secretion [8,14,23], reactive oxygen species (ROS) fluxes [24–26], Ca^2+^ fluxes, mitochondrial polarity, cell wall modification [24,27,28], transportation, MSF drug transportation/ detoxification, and a general fight for trace minerals like iron, magnesium, and copper, including siderophore sequestration [16,29,30]. Attempts to define conserved cellular responses through immediate transcriptional and metabolomic responses to microbial proximity have led to the hypothesis that a conserved fungal immune system likely overlaps with Heterokaryon Incompatibility (HI) responses [15,31,32], physical damage responses, nutrient starvation, cell wall stress responses, antimicrobial responses, and general stress responses [33].

Specifically, *Fusarium graminearum* ROS production, iron sequestration, and antimicrobial genes have been observed as differentially expressed after inoculation of common bacterial microbial associated molecular patterns (MAMPs), including flagellin and peptidoglycan [29]. ROS dynamics and cell-wall modification has been observed in *Rhizopus microsporus* as *Mycetohabitans rhizoxinica* (family Burkholderiaceae) attempt to ingress into the hypha [24]. In fact the T3SS, and resulting transcription activator-like (TAL) effectors function to help the survival of the intracellular *M. rhizoxinica* [34,35] [36]. The general secretory pathway of *Burkholderia gladioli* (BG164) is necessary for cavity disease in *Agaricus bisporus* [37]. The lipopeptide (ralsolamycin) induces chlamydospore formation and Bikaverin production in many fungal species [9,28]. Both *Bacillus subtilis* and *Escherichia coli* induce the same secreted antimicrobial peptides in *Coprinopsis cinerea* [14]. Interestingly, the mitogen activated protein kinase *Tmk1* in *Trichoderma atroviride* not only plays a key role in hyphal regeneration after physical damage [33], but also in the creation of antimicrobial peptides [16] demonstrating the link between signaling networks, antimicrobial and damage responses. In addition, various RNA interference mechanisms recognize viral genomes [10,38]. To reiterate, in all of these cases no upstream non-self recognition system has been linked to MAMP or effector recognition, signaling cascades, metabolite/antibiotic production, or fitness outcomes.

Non-self pattern recognition receptors in fungi, featuring genes governing HI recognition, can be likened to fungal innate immune sensors/receptors, given that recognition of non-self affects resource plundering by invading organisms [17]. Indeed, the fungal non-self recognition field did not hypothesize HI processes as part of, or an evolutionary product of, a “fungal Innate Immune system” until proteins resembling the architecture of NBD-LRR domain-containing (NLR) proteins were discovered to be active in HI in *Podospora anserina* [19,32,39,40] and the cell death reaction was compared to innate immune responses that plants and animals have to colonizing microbes [20,41]. Since the first NLR mechanistic discovery, evolutionary analyses and comparative genomics concluded that although LRR domains are scarce within fungi [42], there exists a diverse repertoire of “NLR-like” proteins amongst the Dikarya with three main repeat domains: WD40, ANKYRIN, and TPR [43,44]. Mechanistic evidence has been reported for four NLR-like proteins in *P. anserina (het-d, het-e, het-r,* and *nwd2)* [39,40,45,46], two NLR-like proteins in *Cryphonectria parasitica (vic2* and *vic4)* [47,48] and one NLR-like protein from *Neurospora crassa* (*plp-1*) [49] mediating the HI allorecognition cell-death reaction, similar to protective cell death in plant, animal, and bacterial innate immune systems.

To investigate the link between microbial recognition, possible immune pathways and host/microbe fitness, we selected *N. crassa* as a study system. *N. crassa* has an active research community, a full deletion collection, genetic manipulation protocols, a complete and annotated genome, and studies of cell death induced by HI and other antimicrobial agents. In fact, *N. crassa* is now a model for fungal virology [50] making it a rising system in fungal-microbial interaction studies. In Wichmann et al (2008), bacterial growth measurements suggest that the bacteria *Pseudomonas syringae pv. tomato DC3000* (PSTDC3000) can utilize *N. crassa* as a sole nutrient source to proliferate, and travel along the mycelium [51]. Using vital dyes, cell death was not observed in hyphae upon exposure to bacteria, suggesting that PSTDC3000 has a “biotrophic” or “semi-biotrophic” interaction with *N. crassa*. *N. crassa* also produces very few antimicrobials, making it an intriguing system to investigate how fungi protect themselves from extracellular bacterial proliferation without an extensive antimicrobial repertoire [52].

In Wichmann et. al (2008) *P. syringae* and *N. crassa* interactions are studied because PSTDC3000 *and P. syringae pv. Syringae B728a* are found to harbor a gene, *phcA*, that is homologous to the *het-c* gene of *N. crassa* [51]. In fact*, phcA* is conserved in many *P. syringae* strains. The *phcA mutant of* PSTDC3000 had a slight fitness defect in colonizing naked hyphae of *N. crassa* in the beginning of the interaction and was shown to initiate HI and form a heterocomplex with *het-c* during ectopic expression in *N. crassa*. The ecological relevance of *N. crassa* existing with *Psuedomonas syringae* species was hypothesized on the basis that both organisms are epiphytic for at least a portion of their lifecycle, but direct evidence did not exist. Likewise, direct evidence of the deliverance of the *phcA* protein naturally from the bacteria to the fungus was not observed. The two organisms were used because of existing genetic resources and organismal understanding, similar to how PSTDC3000 is a natural tomato pathogen but used in the laboratory to infect *Arabidopsis thaliana* and *Nicotiana benthamiana* in order to study conserved portions of the plant immune system [53,54].

Dikarya, like *N. crassa,* are also known to harbor a more numerous and diverse repertoire of predicted NLR-like proteins than early-diverging fungi and Saccharomycotina yeasts [43,55] [49]. Other evidence of a putative immune system in *N. crassa* includes many canonical allorecognition systems, allowing separation between individuals either before, during or after fusion. These loci are: *doc* governing communication at a distance [56], *cwr* which causes arrest and cell wall accumulation upon contact [57] and at least 14 canonical het loci [58,59] that govern compatibility post cellular fusion. Vegetative fusion between germlings/hyphae that contain incompatible *sec-9/plp-1* alleles results in cell death. As previously mentioned, *plp-1* encodes a NLR-like protein with a patatin phospholipase domain [49]. When cells/hyphae that contain incompatible *rcd-1* alleles undergo vegetative cell fusion, the fusion compartment undergoes rapid cell death. The *rcd-1* locus encodes a homolog of Gasdermin [60], which induces a cell death reaction termed pyroptosis in mammalian innate immunity reactions [61]. In particular, co-expression of incompatible *rcd-1* alleles in Human 293T cells triggered a pyroptotic-like cell death [62], suggesting conservation of gasdermin function between fungi and animals.

Here, we report that *Pseudomonas* species co-occur with various *Neurospora* species in natural habitats, including *Pseudomonas* “9.3L” that is phylogenetically related to *P. syringae.* We used direct inoculation of PSTDC3000 onto *N. crassa* hyphae or germlings to establish a “pathosystem”. We observed environmental, time, and tissue-dependent fitness of both *P. syringae* and *N. crassa* when interacting directly or at a distance. Using flow cytometry of germlings, we measured changes in cell wall integrity in a subpopulation of fungal cells as early as ten minutes after contact with bacteria. We also captured the early transcriptional time points after bacterial inoculation onto fungal tissue that allowed us to observe possible cellular processes protecting *N. crassa* or being manipulated by bacteria. Using *N. crassa* deletion mutants in categories identified by transcriptional profiling, we assessed cell death and overall fitness phenotypes of *N. crassa* when challenged by bacteria. This study provides a tractable, biosafe “pathosystem” that treats fungi as hosts to extracellular bacterial colonization with higher throughput methodologies to study immune outputs, signalling cascades, cell death, general fungal fitness, and bacterial colonization.

## Results

### *Neurospora* co-exists with several *Pseudomonas* species in native ecological environments

We sought out direct evidence that *Pseudomonas* species were co-occurring and colonizing *Neurospora* environmental isolates by sampling two wildfire-affected sites in Southern France: Villeveyrac (V) and La Grande Motte (L), approximately 50 kilometers apart. All 87 *Neurospora* samples were grown directly from the wood chips onto R2A agar without added antibiotics, allowing direct observation of extra-hyphal bacteria traversing along the hypha of *Neurospora* (**Figure 1A**). Approximately twenty-five percent (22 out of 87) of the samples grew out with extra-hyphal bacteria traversing along the hypha **(Figure 1B)**. Samples from Villeveyrac had a much smaller ratio of hyphal bacteria (9/66); while La Grande Motte had bacteria on a majority of the samples (13/21). Of the total bacteria growing out, three strains from Villeveyrac and two from La Grande Motte were *Pseudomonas sp*. **(Figure 1C)**. The *Pseudomonas* isolated from the *Neurospora* sample 9.3L was closely related to *P. syringae* **(Figure 1C)**. These data indicate that wild *Neurospora sp.* isolates interact with bacteria, including *Pseudomonas sp*., and that bacteria associated with *Neurospora* can traverse the hyphae in R2A media.

**Figure 1:**
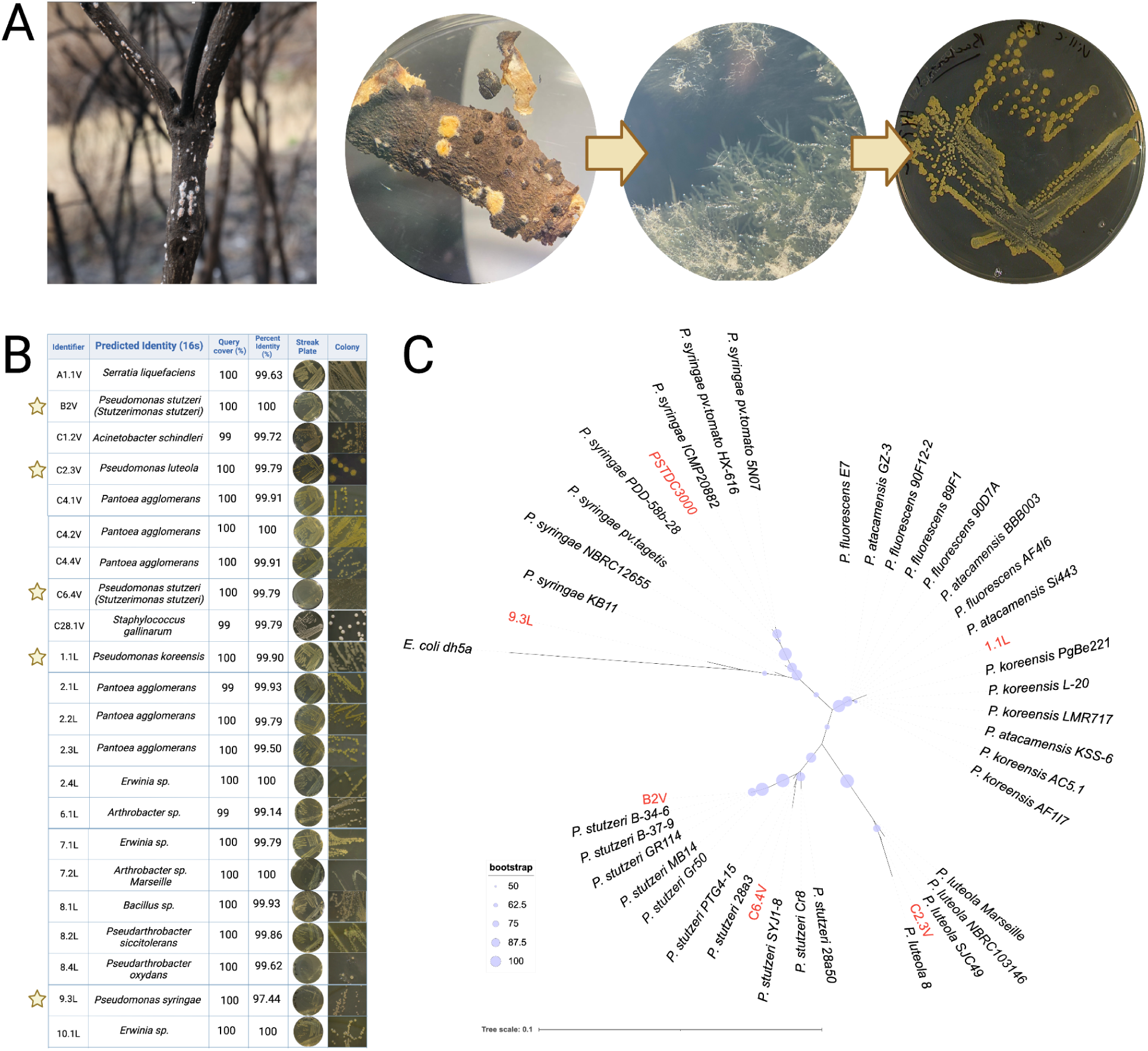
*Pseudomonas sp.* interact with *Neurospora* species in the environment, and traverse hyphae. A) Photo of *N. crassa* conidiation after fire on Scotch broom (*Cytisus scoparius*) in Villeveyrac, alongside sampling protocol from burnt wood chips containing visible *Neurospora* conidiation to obtain bacteria. B) Twenty-two samples grew out with bacteria present along the hyphae: five of which were *Pseudomonas* species (yellow star). C) Maximum likelihood phylogeny of the 16s region shows relation of the five environmental samples of *Pseudomonas* species to *Pseudomonas syringae DC3000* and other reference strains.

### Interaction between PSTDC3000 and *N. crassa* alters bacterial and fungal growth depending on environment and tissue

Having demonstrated that *N. crassa* and *P. syringae* co-occur in nature, we focused on understanding the molecular basis for interactions of the model system PSTDC3000 and *N. crassa*. We focused on PSTDC3000 due to availability of genetic resources and its well-characterized interactions with plant immune systems. First, we compared how fungal cells responded to PSTDC3000 at a distance. Exposure of *N. crassa* to PSTDC3000 and *E. coli* DH5a at a one-inch distance, applied in 10ul spots on the same agar plate, resulted in an alteration of fungal growth kinetics that was dependent on the dose of bacteria and growth media, suggesting that fungi can sense bacterial presence before direct contact **(Supplementary** Figure 1). Nutrient poor Vogel’s Minimal Medium (VMM) and nutrient rich Luria Agar (LA) had opposite effects on growth kinetics of *N. crassa* when challenged with bacteria **(Supplementary** Figure 1A). *N. crassa* has increased conidiation on VMM seeded with bacteria **(Supplementary** Figure 1B) while *N. crassa* has decreased conidiation on LA seeded with bacteria. This distance related interaction seems mostly affected by diffusible molecules and less by volatile **(Supplementary** Figure 1C), supporting the role of long-distance diffusible molecules in directing interaction outcomes.

An inoculation-based approach consisting of putting bacteria directly onto *N. crassa* tissue at different growth stages revealed the cellular reaction to bacterial immediate proximity (**Figure 2A**). Immediately, upon being inoculated onto the fungal cells, PSTDC3000 preferentially surrounded *N. crassa* conidia, germlings, and hyphae rather than the environment **(Figure 2B)**. However, this was also observed for interaction between PSTDC3000 with glass beads, and fiberglass **(Figure 2B)**, and we observed similar phenotypes for *E. coli* DH5a **(Supplementary** Figure 2A). This result suggested the clustering of bacteria naturally occurs on any solid surface upon inoculum evaporation, as reported on before [63], and is not solely due to a biological process such as chemotaxis. These experiments suggest that forced proximity may be a useful protocol in standardizing extra-hyphal bacterial interaction assays. Furthermore, condensation of bacteria onto solid surfaces is likely ecologically-relevant, as *Pseudomonas* species exist within precipitation [64–66].

**Figure 2:**
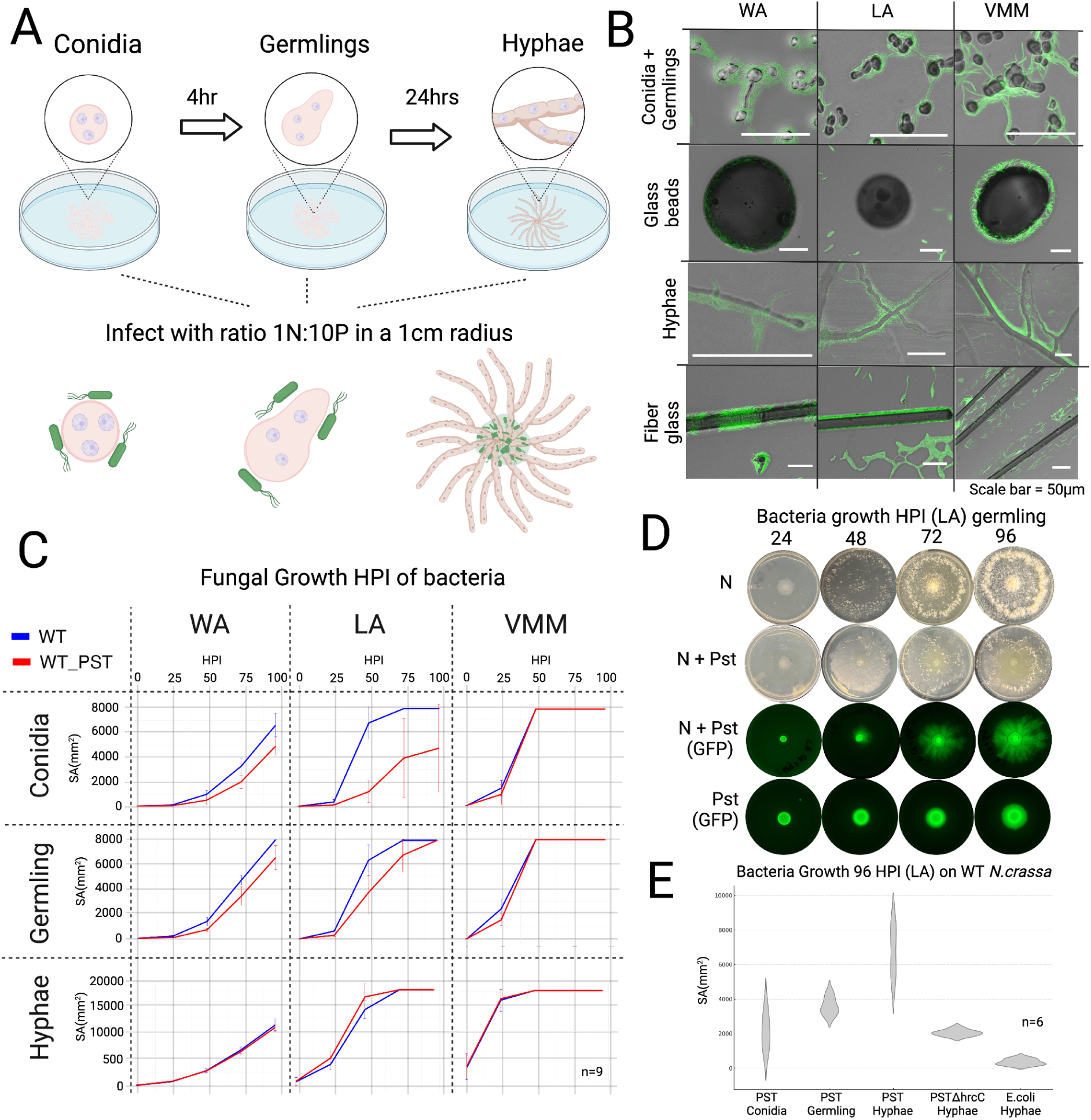
Inoculation of *Pseudomonas syringae* PSTDC3000 directly onto *Neurospora crassa* conidia, germlings, and hyphae causes environmental/tissue dependent growth response and bacterial proliferation. A) *Neurospora crassa* OR74A tissue types and protocols of PSTDC3000 inoculation used in study. B) Inoculation of bacteria directly onto *N. crassa* forces bacteria into the hydrosphere. C) Surface Area (SA) growth curves of fungal tissue with and without PSTDC3000 exposure over time showing inoculation on conidia and LA having the most defined fitness defects. D) Spread of bacteria is easily measured and observed in LA, showing various fitness differences within the co-culture. E) Bacterial spread is inversely correlated with *N. crassa* development, showing fastest spread after inoculation on pre-established hyphae (n=6). The type three secretion mutant (ΔhrcC) of PSTDC3000 more slowly spreads over the hyphae whereas *E. coli* cannot swim along *N. crassa* hyphae. The data used in this figure can be found in **Supplementary Table 1**.

To define the bacterial-fungal interactions, we measured resulting fungal fitness and bacterial proliferation upon interaction in three different environments and three separate tissue types **(Figure 2C).** Inoculating PSTDC3000 on top of conidia, germlings or mature hyphae reduced growth of *N.crassa* in all environmental conditions early in the interaction **(Figure 2C)**. Fungal growth and bacterial proliferation on hyphae measured over time revealed that growth defects were most drastic in LA and for conidia. *N. crassa* colonies growing in water agar (WA) showed some fitness defects at ∼48 hrs, whereas colonies growing in VMM showed the greatest growth defect at 24 hrs. Spread of bacteria was only measurable in LA. No bacteria were observed along the hyphae in WA and VMM **(Figure 2D)**, even though PSTDC3000 was able to rapidly swim in all environments on fiberglass **(Supplementary** Figure 2B). Bacterial spread is inversely correlated with *N. crassa* vegetative development, showing fastest spread after inoculation on pre-established hyphae **(Figure 2E)**. Interestingly, the type-three secretion mutant (ΔhrcC) of PSTDC3000 more slowly spreads over the hyphae **(Figure 2E).** We recapitulate the result from Wichmann et. al (2008) showing that *E. coli* cannot swim along *N. crassa* hyphae, but this time in the LA environment **(Figure 2E)**.

### PSTDC3000 induces Propidium Iodide uptake in a subpopulation of *N. crassa* germlings as early as 10 minutes post-contact

We investigated the earliest time points of the interaction since innate immune receptors can respond to bacterial inoculation as early as 10 minutes after exposure, as reported in other systems [67,68]. HI reactions governed by a NLR-like protein *plp-1* in *N. crassa* are also initiated rapidly at around 20 minutes post fusion of germlings and commonly involves a programmed cell death reaction [49]. To observe the earliest time points in fungal response to coming in contact with bacteria, we performed flow cytometry on germlings inoculated with PSTDC3000 and measured if programmed cell death was occurring by uptake of the vital dye Propidium Iodide (PI) **(Figure 3A).** The flow cytometry protocol was adapted from the germling specific protocol in Heller *et. al* 2018 [49] that assessed HI induced cell death using propidium iodide uptake also in *N. crassa* germlings.

**Figure 3:**
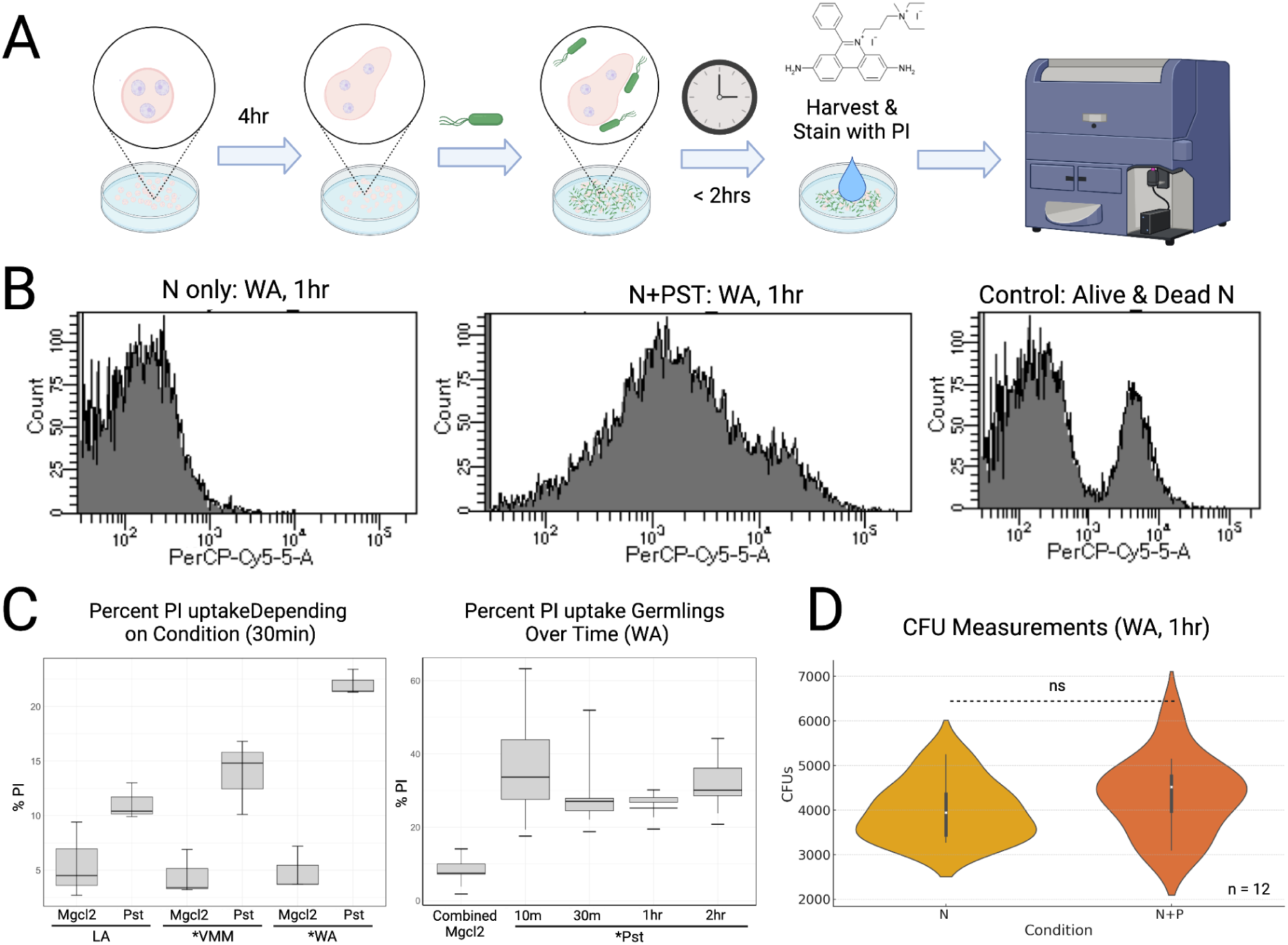
Cell death assays using vital dyes and colony-forming unit (CFU) counting after PSTDC3000 inoculation reveals Propidium Iodide (PI) uptake but no cell death happening in the early timepoints of inoculation. A) Simplified protocol of inoculating *N. crassa* germlings with bacteria, staining with vital dye, PI, and imaging them on the flow cytometer. B) Flow cytometry graphs showing the distribution of uptake of PI in ∼20,000 *N. crassa* germlings in the presence and absence of PSTDC3000. The negative control was a 50% mix of living conidia and conidia killed by heating. C) The percentage of *N. crassa* cells stained with PI in the population is environmentally dependent and steady over time (< 2hrs), showing PI-stained cells as early as 10 minutes. D) CFU measurements of *N. crassa* in the early timepoints verified that true cell death did not occur (P value = 0.28. Anova P value cutoff = *0.01). The data used in this figure can be found in **Supplementary Table 2**. The exact protocol for flow cytometry and CFU countings of *N. crassa* germlings is found in **Supplementary File 1** and **Supplementary File 2**, respectively.

Imaging ∼20,000 germling cells of *N. crassa* with PI reveals an uptake of PI fluorescence in the presence of PSTDC3000. However, a bimodal distribution of mixed living and dead cells was rarely observed, suggesting only a slight PI uptake **(Figure 3B)**. PI is known to enter non-viable cells through cell wall/membrane damage, but has also been known to enter viable fungal cells during immediate exposure to non-lethal cell wall/membrane stress and fluorescences slightly more dim than cells that are truly dead [69]. This indicates membrane stress induction and then repair [69,70]. The intermediate phenotype suggested that programmed cell death was not occurring **(Figure 3B).** The percentage of *N. crassa* cells taking up PI in the population was environmentally dependent and steady over time (< 2 hrs), showing PI-stained cells as early as 10 minutes post bacterial inoculation and the highest percentage of PI-stained cells in WA environment **(Figure 3C)**. Colony-forming unit (CFU) measurements of *N. crassa* in the early time points using 2% sorbose verified that cell death was not occurring under these conditions as the number of *N. crassa* colony forming units were statistically similar with and without PSTDC3000 treatment **(Figure 3D).**

### RNA-seq of *N. crassa* germlings upon contact with PSTDC3000 reveals time-dependent transcriptional response to bacteria

To get a view of the fungal transcriptional response occurring at the earliest time points, we performed an RNA-seq experiment at 10 minutes and 1 hour post PSTDC3000 inoculation of *N. crassa* germlings using the WA environment due to the greatest percentage of cells with PI uptake. Principal component analysis revealed that two principal components (PC) account for 33% and 15% of transcriptional variance, corresponding to pathogen exposure and time, respectively. This result verified that transcriptional change was rapid and induced by the presence of bacteria **(Figure 4A)**. In total, 1,863 genes (∼17% of genome) were significantly differentially expressed (DE) during the two time points after exposure of bacteria by selecting a fold change (Log_2_FC) ≥ 1, and (Log_2_FC) ≤ −1, false discovery rate (FDR) < 0.05. Between the time points,157 up-regulated genes were shared, 297 genes specific to 10 min, and 346 specific to 1 hr post bacterial inoculation **(Figure 4A)**. 253 down-regulated DE genes were shared, leaving 391 genes specific to 10 min and 413 to 1 hour **(Figure 4B)**. Interestingly, only five genes were downregulated at 10 min and upregulated at 1hr highlighting up-regulated and down-regulated genes as distinct.

**Figure 4:**
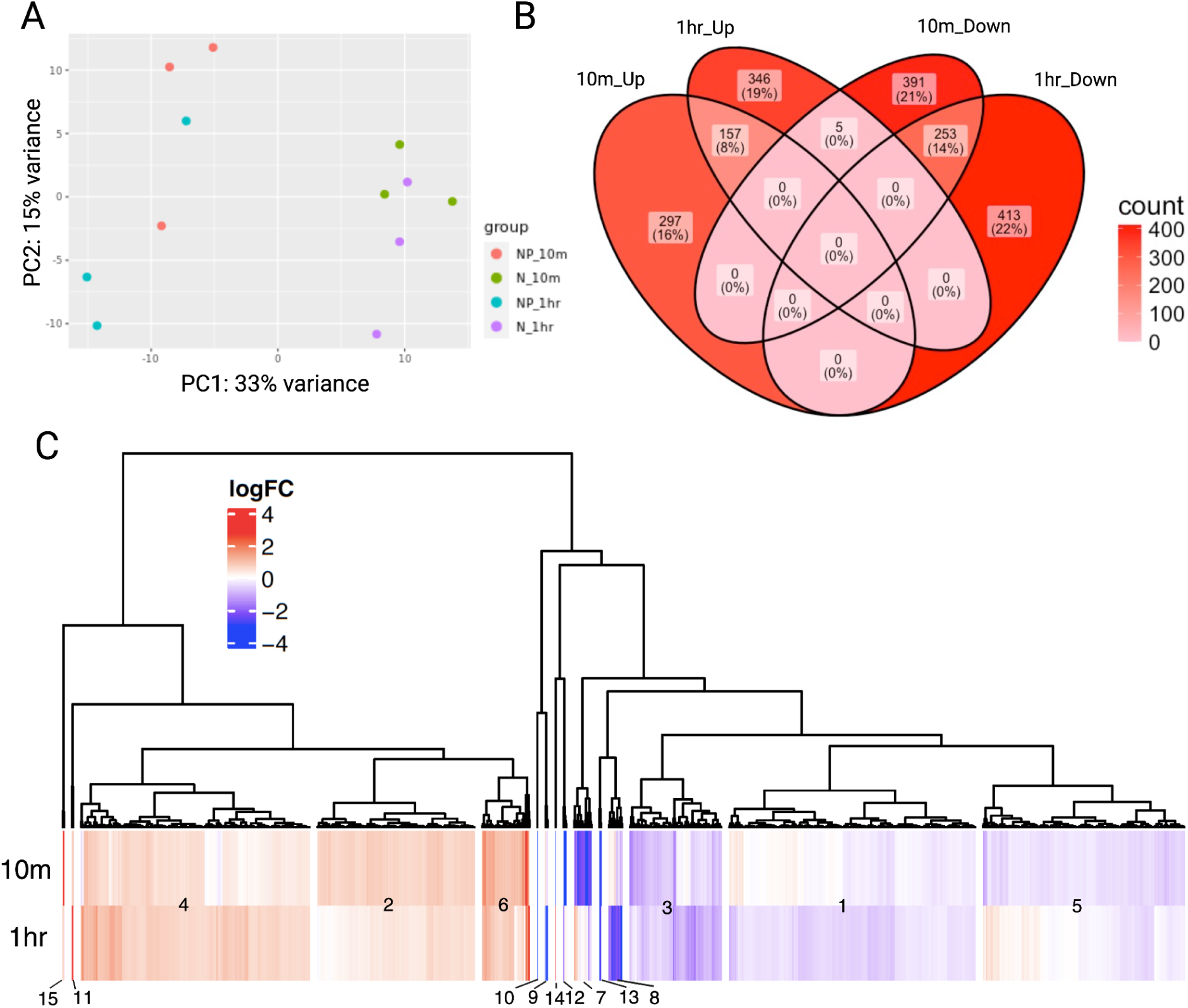
RNA sequencing of *N. crassa* germlings on 10minute and 1hr bacteria exposure reveals transcriptional response. A) Principal component analysis of RNA sequencing samples after normalization. “N” refers to *N. crassa* inoculated with MgCl2 while “NP” refers to *N. crassa* inoculated with PSTDC3000 in MgCl2. “10m” refers to the ten minute time point while “1hr” refers to the one hour time point. B) Venn diagram of number of genes shared between time points highlights up-regulated and down-regulated genes as distinct C) Hierarchical-clustering of the heat map of all genes differentially expressed in this assay and split into 15 clusters based on normalized Log2FC values and time. The data used in this figure can be found in **Supplementary Table 3.**

### FunCat, GO and KEGG enrichment analysis

We next performed hierarchical clustering of all DE genes based on time and the Log_2_FC upon PSTDC3000 inoculation and identified 15 clusters (Figure 4C). We then grouped the genes within each cluster to shared, 10 min-specific and 1hr-specific timepoints (**Table 1**). We then examined the enrichment of functional annotations within each cluster by performing FunCat [71], GO [72], and KEGG [73] enrichment analysis using the tool FungiFun [74]. A detailed supplementary file on the enrichment can be found in **Supplementary Table 3.**

**Table 1:** Summary of FunCat, GO and KEGG enrichment analysis on each cluster. The full enrichment analysis can be found in Supplementary Table 3 and a description of each cluster can be found in Supplementary File 3.

|  | **10min** | **Shared** | **1hr** |
| --- | --- | --- | --- |
| **Log<sub>2</sub>FC**<br>**(1-10)** | Cluster 2<br>•rRNA processing,<br>•ribosome biogenesis<br>•Ribosomal proteins<br>•RNA polymerase<br>RNA binding<br>DNA directed RNA<br>polymerase activity<br>•Transcription<br>•Translation<br>•Intracellular | Cluster 6<br>•Cell wall dynamics:<br>polysaccharide metabolism (**lyz**)<br>•**Heavy metal ion**<br>**transmembrane transport** (Cu+,<br>Fe3+, etc.) (**tcu-1, fer-1**)<br>•**Sod-2, mdr-6**<br>•Pyridoxal phosphate binding<br>•Citrate cycle, Krebs cycle, TCA<br>cycle<br>•Mating pheromone activity<br>matA-1 (NCU01958)<br>•Nitrogen, carbohydrate<br>metabolism | Cluster 4<br>•Protein folding & stabilization<br>(Thioredoxin-2 & 4, ROS, apoptosis)<br>Heat Shock Proteins<br>•mitotic chromosome condensation<br>(condesin complex)<br>•protein processing in the ER<br>•Cell wall dynamics: polysaccharide<br>metabolism (Glycoside hydrolase) |
| **Log<sub>2</sub>FC**<br>**(>10)** | Cluster15<br>•Ubiquitin mediated<br>proteolysis<br>**(NCU09866T3) E3**<br>**ubiquitin-protein**<br>**transferase**<br>(NCU01000T0) ddp-9<br>dipeptidyl peptidase |  | Cluster 11<br>•Tubulin dependent transport<br>(NCU05180T0) kin-9<br>•**Leashin-woronin body tether**<br>**(NCU02793T3)**<br>•bdp-3 (NCU08423T0) |
| **Log<sub>2</sub>FC**<br>**(-4) -(-1)** | Cluster 5<br>•C-compound and<br>carbohydrate<br>metabolism<br>•Lipid/fatty acid<br>metabolism & transport<br>•Glutathione<br>conjugation reaction<br>•Superoxide<br>metabolism/ oxidation<br>reduction<br>•NADP binding<br>•Extracellular<br>region/peroxisome<br>•Biosynthesis of<br>secondary metabolites<br>•Autophagy/ early<br>endosome | Cluster 3<br>•Nitrogen, selenium, sulfur<br>metabolism<br>•Amino Acid metabolisms<br>•Transport Facilities<br>•Import, ABC, siderophore,<br>sulfate, ion, C-compound and<br>carbohydrate transport<br>•oxidative stress response | Cluster 1<br>•Assimilation of ammonia<br>•Sulfur metabolism<br>Glycerophospholipid metabolism<br>•C-compound and carbohydrate<br>metabolism<br>•NLR-like transcript, *adv-1* and sexual<br>development proteins/pheromones<br>•Integral component of membrane |
| **Log<sub>2</sub>FC**<br>**(-15) -(-4)** | Cluster 7<br>•Meiotic recombination<br>•MAPK<br>signaling/mating-type<br>pheromone receptor<br>**activity (pre-2) GCPR** | Cluster 13<br>•Ubiquitin mediated proteolysis<br>NCU09866T2<br>•Drug transmembrane transport | Cluster 8<br>•Glycerophospholipid biosynthetic<br>process |
| **Log<sub>2</sub>FC<**<br>**(-15)** | Cluster 12<br>•Includes **NLR-like**<br>**transcript (NCU09760)**<br>and *sec-7*<br>•Regulation of<br>transcription<br>•DNA binding<br>•Chromatin<br>modification<br>•GTPase ARF protein<br>signal transduction | Cluster 10<br>•Woronin body tether<br>**NCU02793T2** | Cluster 9<br>•Ubiquitin mediated proteolysis<br>**NCU09866T1**<br>•MAPK signaling NCU02234 (*mik-1*) |

This analysis revealed a wide range of biological processes as being attuned after bacterial inoculation. Including transcription, translation, primary metabolism, secondary metabolism, amino acid metabolism, nitrogen metabolism, lipid metabolism, and carbohydrate metabolism, with enrichment for cell wall-modifying enzymes and secreted proteins. Many expected processes of “cell stress” pathways and detoxifying mechanisms were seen including oxidative stress responses, mitochondrial polarity, a glutathione conjugation reaction, protein degradation regulation, secreted proteins, transport facilities, import, nonvascular import, ABC transporters, MFS drug transporters and genes implicated in trace-metal warfare. There was evidence of the CWI pathway being down regulated because of the presence of mik-1, pre-2 and adv-1. Interestingly, mating pheromone activity and the mating type loci appeared in this data set. A more detailed analysis of each cluster can be found in **Supplementary File 3**.

### Testing *N. crassa* mutants informed by RNA-seq reveals phenotypic variation in fitness after exposure to PSTDC3000

We selected fourteen initial mutants of *N. crassa* from both the upregulated and downregulated gene sets that may be involved in defense and defense signalling. We then observed if their disruption affected fitness outcomes in Neurospora and PSTDC3000 growth **(Figure 5)**.

**Figure 5:**
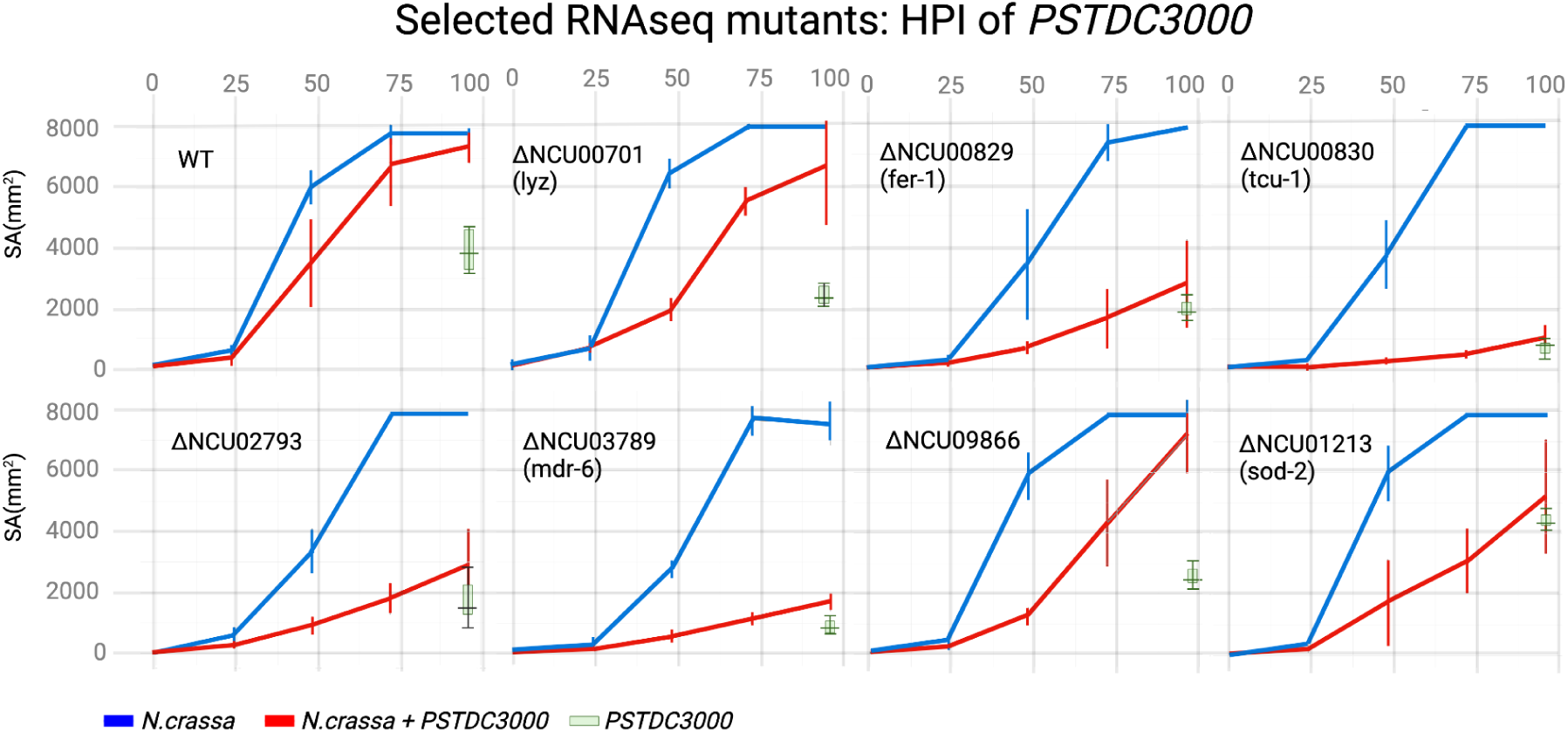
Growth curves for mutants selected from RNAseq analysis reveals various phenotypic defects. A) Growth curves of 7 mutants of *N. crassa* that showed significant differences in growth as compared to wildtype after PSTDC3000 inoculation on LA media (n=9). Significance was assayed using area under the curve (AUC) analysis coupled with paired T-tests.The data used in this figure can be found in Supplementary Table 4.

Within the shared upregulated gene set (cluster 6), we used mutants in all three genes associated with the “Heavy Metal Ion transport” functional category. Both of the neighboring transcripts: NCU00829 (ferric reductase; *fer-1*) and NCU00830 (copper transporter; *tcu-1*) showed the most extreme fitness defects in vegetative growth after PSTDC3000 pressure compared to uninoculated controls **(Figure 5)**. Interestingly, NCU003281 (copper transporter; *tcu-2*) showed slightly higher growth when paired with PSTDC3000 colonization **(supplementary Figure 3).** Close in this cluster included NCU01213 (superoxide dismutase; *sod-2*) which also showed a strong fitness defect with bacterial inoculations **(Figure 5)**. Within this cluster was also a secreted lysozyme - glycoside hydrolase (*lyz*; NCU00701) that showed homology to lysozymes produced by *Coprinopsis cinerea* in response to bacteria [14]. The mutant for this lysozyme only showed a slight fitness defect in the presence of bacteria **(Figure 5).** The highly upregulated gene NCU03788 (cysteine synthase B; *cys-19*) and neighboring transcript NCU03789 (multi-drug resistance-6 protein; *mdr-6*) were also tested. While no fitness difference was observed for *cys-19* **(Supplementary** Figure 3), a large fitness defect was observed for *mdr-6* **(Figure 5)**.

We then selected a few genes based upon their expression pattern and functional annotation encoding well-studied proteins. NCU02793, the most downregulated gene in both timepoints, encodes for the functionally characterized leashin Woronin-body tethers: *lah-1* and *lah-2* [75] which locate the Woronin-bodies, fungal organelles that seal septal pores, to either the septal pore rims or randomly along the inner membrane of *N. crassa* [75]. The mutant of NCU02793 also showed significant fitness decreases with PSTDC3000 colonization **(Figure 5).** From the list of genes upregulated at 1 hour, only two sugar transporters were tested: NCU05853 (sugar transporter-12; *sut-12*) and NCU08114 (hexose transporter; *cdt-2*). Both showed very similar growth phenotypes to wildtype *N. crassa* when exposed to bacteria **(Supplementary** Figure 3).

We also selected a few genes from the receptor dataset as possible receptors of bacterial ligands, or executors of the cell death pathway. One NLR gene (NCU09760) out of the ∼19 encoded in the *N. crassa* genome was extremely downregulated at 10min **(Supplemental Table 3)**. This mutant showed little to no fitness defect in the presence of PSTDC3000 **(Supplementary** Figure 3). The HECT-type E3 ubiquitin transferase “thyroid hormone receptor interactor” NCU09866; which had one transcript upregulated, and two downregulated, showed a slight fitness defect **(Figure 5)**.

## Discussion

It is hypothesized that all non-self pattern recognition receptors in fungi governing recognition of non-self (such as HI) can be likened to a fungal immune system, given the fact that the processes stop resource plundering from non-self [17]. However, no intracellular or extracellular receptor recognizing microbial-specific ligands have been identified besides adenylyl cyclase Cyr1p in *Candida albicans* which recognizes peptidoglycan [22]. Here we devised a model pathosystem of *N. crassa and* PSTDC3000 to begin testing hypotheses of the fungal immune system. We first showed that *N. crassa* naturally exists with bacterial pressure in post fire environments. We then set up fitness assays measuring PSTDC3000 and *N. crassa* growth after direct contact, demonstrating that the outcome of their interactions is context dependent, tissue dependent, and time dependent. Use of the RNA-seq data set at early time points in the interaction allowed us systematic testing of various mutants that could be contributing to the survival of *N. crassa* that are rapidly induced in the presence of bacteria. We then categorized these genes as possible receptors, signaling systems and a putative “immune” response.

Here, we report ctr copper transporter (*tcu-1*), ferric reductase (*fer-1*), superoxide reductase (*sod-2*), multidrug resistance transporter (*mdr-6*), a secreted lysozyme-Glycoside hydrolase (*lyz*), and the Woronin body tether (NCU02793), as being both differentially expressed and functionally necessary for the survival of *N. crassa* under bacterial pressures **(Figure 5)**. **Figure 6** is a graphical summary of what we found in this paper.

**Figure 6:**
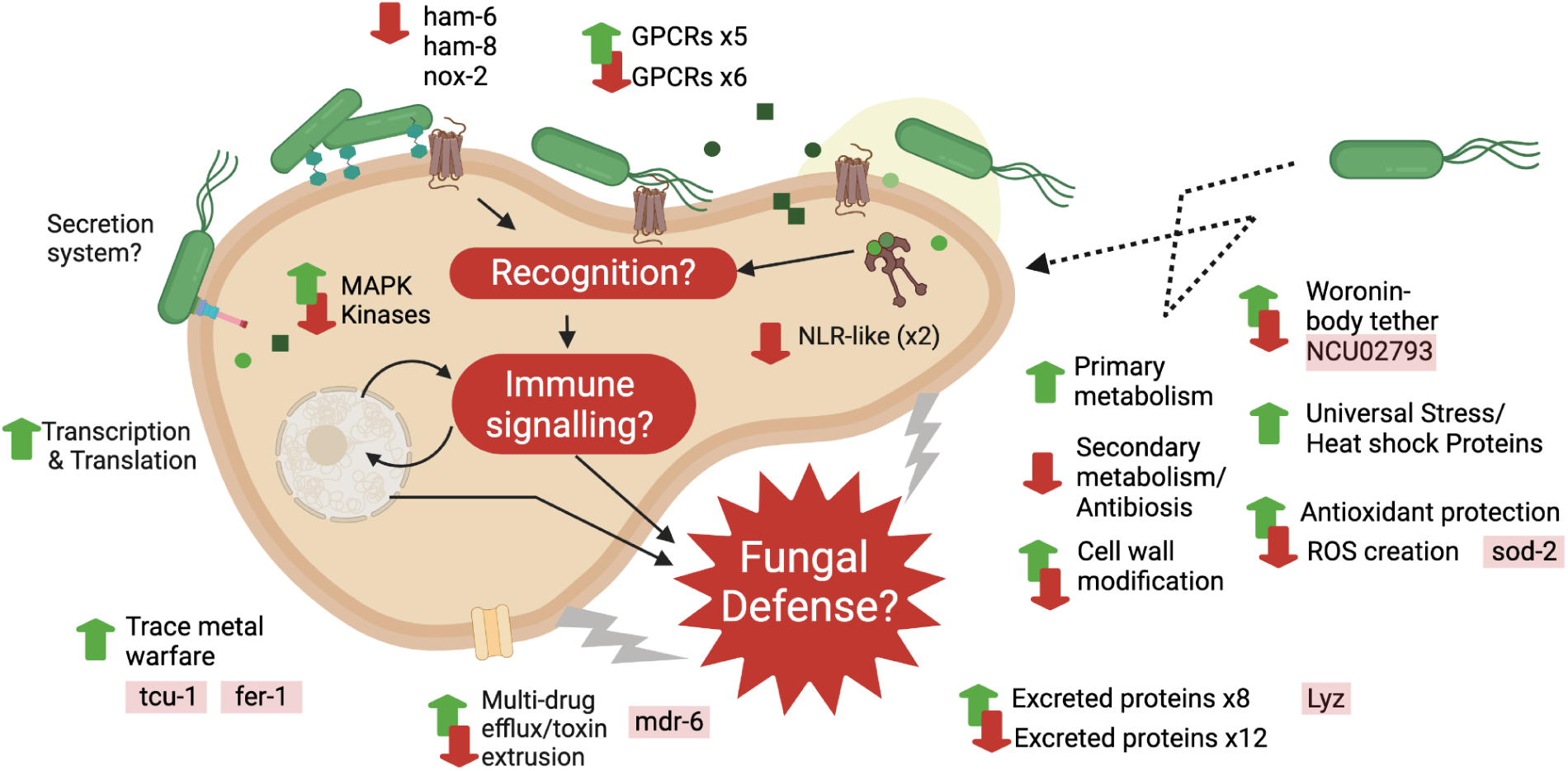
Summary of *Neurospora crassa*’s predicted interaction to PSTDC3000. Upregulated processes are shown as green up arrows where downregulated processes are shown as red down arrows. Genes identified as being important to the survival of *N. crassa* under PSTDC3000 pressure are highlighted in red and placed under the category they are hypothesized to be involved in.

Modulation of ROS (*sod-2*), secreted proteins (*lyz*), antimicrobial synthesis, and detoxifying mechanisms (mdr-6) are a common defense mechanism seen in fungal-bacterial interactions [5,9,14,15] as well as human-bacteria and plant-bacteria interactions. Linking how these systems are induced or manipulated by bacteria will surely address conserved portions of the fungal immune system. Likewise, *lyz* is analogous to the antibiotic Glycoside Hydrolase Lysozyme in *Coprinopsis cinerea* induced by both *Bacillus subtilis* and *Escherichia coli* [14]. Further work will be done on how this gene is attuned and conserved across fungal-bacterial interactions. Additionally, it would be of interest to see if any of the secreted proteins have antimicrobial activity once purified.

Likewise, copper, iron and biotin availability have been shown to modulate a wide range of fungal/bacterial interactions as a mechanism of controlling microbial growth [5,16]. Trace metal warfare is also medically relevant in modulating pathogen growth in mammals [76,77]. The membrane transporter tcu-1 has also been shown to be transcriptionally stimulated by trace metal starvation [78]. Previously, NCU00829 has been shown to contribute to cellulase activity/efficiency by producing hydroxyl radicals from fenton chemistry [79]. Membrane-associated ferric reductase activity has also been shown to reduce insoluble Fe3+ to Fe2+ and then the membrane transporter delivers the ferrous ions inside the cell. The reductase and the transporter are known to be controlled by availability of ions, suggesting that the two genes are expressed by iron or copper starvation induced by PSTDC3000.

NCU02793, the leashin Woronin body tether, showed up as the one of the most downregulated (NCU02793T2) genes in both timepoints. However, a separate transcript showed up as upregulated at the one hour time point (NCU02793T3). Regardless, the mutant spanning the entire gene (NCU02793) showed a major defect in survival against PSTDC3000. The tether is known to locate the Woronin bodies either to the septal pore rims or randomly along the inner membrane of *N. crassa*. Two transcripts have been functionally analyzed from this locus: *Lah-1* and *lah-2* [75] Loss of *lah-1* is known to greatly increase hyphal “bleeding” after wounding and tether specifically to woronin bodies, while lah-2 localizes to septal pores. We hypothesize that the function of the tether in this interaction is to assist in wound repair of the cell-membrane, balancing cell damage that we observed with PI uptake and preventing cell death. There are now five proposed transcripts for the leashin gene. A more rigorous study of the separate transcripts that are differentially expressed in this dataset will be conducted in the future.

The CFU assays suggest that true cell death is not occurring in the population before one hour, but the responses that have lasting impact on the fitness of *N. crassa* are rapidly induced. *N. crassa* regained its normal growth rate on water agar suggesting that the cells we sequenced were initiating a response that would end in organismal survival and adaptation to presence of bacteria, suppressing regulated cell death. Likewise, we see two NLR-like genes downregulated alongside hyphal anastomosis mutant genes (ham) genes, including evidence of modulation of the Cell Wall Integrity (CWI) pathway (*nox-2*, *mik-1*, *pre-2* and *adv-1*). This survival initiation is similarly seen in the interaction that *Podospora anserina* has with *serratia* species, as NLR-like proteins are not upregulated, unlike during the HI reaction that favors cell death [15]. Similar to the transcriptional responses initiated by HI and PHS in *N. crassa*, we are likely observing both a cell damage that could potentially lead to cell death and survival response in the early time points [31,80]. Whether regulated cell death in sub-population of cells, a mark of immune response in other systems, can be fully executed and would protect fungi from microbial proliferation is still undetermined. Cell death as a protective mechanism may be more relevant for intracellular, biotrophic bacteria or viruses that the fungus can sequester and kill in one hyphal compartment protecting the rest of the colony.

Lastly, within this system it is difficult to distinguish active transcriptional response of *N. crassa* to presence of bacteria from bacteria actively manipulating fungal cells via Type III secretion or other types of secreted proteins. Future work will need to dive deeper into characterizing bacterial response, as we found that the ΔT3SS isolate does not colonize *N. crassa* as well as the wild-type **(Figure 2E)**.

Our study provides a tractable pathosystem, important standardization of protocols, a look into the rapid response *N. crassa* has to the presence of the antagonist PSTDC3000, and some key proteins important in the interaction that are rapidly induced, as summarized in **Figure 6**. Furthermore, many other proteins and processes were observed to be differentially regulated that are yet to be functionally tested, such as G-coupled protein receptors (GCPR), kinases/phosphatases (including *mik-1* and *os-5*), transcription factors, cell-wall modification and attenuation of secondary metabolism. Genes we found most interesting to test in the future are found in **Supplemental Table 3.** Future work will focus on receptors and signaling pathways *N. crassa* uses to respond to PSTDC3000, hopefully uncovering conserved and important networks of the fungal immune system.

## Methods

### Neurospora/bacterial strains, growth, storage and preparation

This study used *Neurospora crassa* OR47A, and environmentally isolated *Neurospora* species. Storage and protocols were done following the <u>FGSC</u> guidelines including creation of various media components. Individual gene knockouts were ordered from the full genome deletion collection hosted by FGSC and stored in 25% glycerol at −80c. Sources of bacteria in this study includes: *Pseudomonas syringae* pv. tomato DC3000, *Pseudomonas syringae* pv. tomato DC3000 GFP [81,82], *P. syringae* pv. tomato DC3000 hrcC-[82] and *E. coli* DH5a.

### Bacterial culturing from *N. crassa* tissue

A small area, that consisted of hyphal cells and conidia, was swabbed from wood chips containing *Neurospora* colonization and placed onto the center of an R2A plate without antibiotics. *Neurospora* and bacterial growth was monitored for 5 days. Bacteria growing out of the hypha was restreaked on a fresh R2A agar plate containing 4 mg/ml amphotericin B until single colonies with distinct morphologies were present. Bacteria from single colonies and representing single species were grown up in liquid culture and stored in −80°C freezer in 60% glycerol.

### Sequence amplification of 16S ribosomal gene and phylogeny construction

A small sample of cells from a single colony was selected for each bacterial plate and suspended in the extraction buffer (RED Extract-N-Amp Tissue PCR Kit,Sigma-Aldrich, cat no XNAT2). The protocol supplied with the kit was then followed to extract DNA and perform PCR. Ribosomal 16s V1-V9 primers were used and both the reverse and forward reads were sequenced through Sanger sequencing to make a consensus sequence by alignment of the full 16s for each bacteria. The 16S consensus sequences for this study can be found on GenBank using the accession number **SUB15006077**. A quick Blast search was performed on each 16s to get to genus level and percent identity and percent query cover was used to assign a tentative species [83]. The Pseudomonas 16s ribosomal Phylogeny was created by downloading reference 16s sequences of selected *Pseudomonas* species from NCBI. Multiple Sequence alignments of reference strains and new sequences of Pseudomonas species was performed using Mafft [84,85], enabling iterative refinement (1000 cycles) and using the L-INS-i algorithm for high-accuracy local pairwise alignment. RAxML-NG [86] was used to create a maximum likelihood phylogeny with default parameters and 100 bootstraps. The phylogeny was visualized and annotated in the Interactive Tree of Life (iTOL) [87].

### Distance co-culture, diffusible and volatile plate assays

Five 10ul drops of an overnight culture of bacteria, adjusted to an OD600= 0.5, was placed in a line half of an inch from the middle of the plate. The bacteria was allowed to grow for a range of days. *N. crassa* OR74A was grown on a VMM plate for two days. We then took five, one-cm diameter agar-plugs of mycelial tips and placed them in a line, half an inch from the middle of the plate, directly opposite the bacteria spots. We then observed growth of *N. crassa* over at least two days. Diffusible and volatile assays began by growing the line of bacteria and then cutting out the bacteria from the agar completely. The resulting agar plate was used to assay growth defects of “diffusible” molecules left by the bacteria on *N.crassa*. The bacterial cells that were cut out of the plate for the “diffusible molecule” assay were used in the volatile assay by placing the bacteria in a “volatile chamber” (a larger plate) that also contained a fresh LA agar plate to grow *N. crassa*. The resulting growth of *N.crassa* was therefore only influenced by volatile molecules (**Supplemental Figure 1).**

### Direct bacterial-fungal interaction assay for growth curves

*N. crassa* OR74A was grown in a glass test tube for 5-7 days until significant macro-conidiation had occurred. Macroconidia was collected from the tube by suspending in 1x PBS and siphoning the suspension through a cheesecloth to remove hyphal fragments. The concentration of conidia was diluted to 4.8 x 10^6 of Neurospora cells per µl of PBS by measuring the optical density (OD) at 600nm. 10µl of conidia was plated on the middle of the plate in a droplet and left to dry. Plates grown for 24 hours were used for hyphal growth curves. Plates grown for 4 hours (at 30**°**C) were used for germling growth curves. Plates used immediately were used for the conidial growth curves. An overnight bacteria culture was collected around an OD600 of 0.8-1 and diluted to 5 × 10^7^ /ml (OD = 0.5) and then washed and resuspended in 10 mM MgCl_2_. 10µl of the bacterial suspension was then inoculated onto the middle of each plate directly over the fungal inoculation spot, estimating a ratio of 1 germling to 10 bacteria cells. The overlay was dried with the lid off in the biosafety cabinet. 10 mM of MgCl_2_ buffer was inoculated onto *N. crassa* as the negative control. Radial surface area was outlined every 24 hours and final pictures were analyzed in ImageJ [88] to calculate surface area growth of bacteria and fungus over time. A more detailed protocol is found in the “Protocol for solid surface interaction assay” section of **Supplementary File 2**.

### Flow cytometry

Cultivation for flow cytometry was identical to cultivation in the solid-surface bacterial-fungal interaction assay except only conidia and germlings could be imaged. The whole plate was inoculated at the same defined cell-cell ratios (1N:10PSTDC3000). The flow cytometry protocol was adapted from the germling specific protocol in Heller et. al 2018 [49] that assessed HI induced cell death using Propidium Iodine. 500ul of conidial suspension was spread with a cell spreader onto plates with cellophane overlay, estimating about 2,400,000 conidia/plate. Cellophane was preferred over pluronics as a way to get *N.crassa* off of the solid surface without damage to hyphae that embed into the agar. Conidia were allowed to germinate for 4 hours at 30**°**C before 500ul of 5 × 10^7^ CFU (OD = 0.5) PSTDC3000 was spread on top and left to dry. After allowing the cells to interact <2hrs, 5ml of 15mM EDTA (pH= 6.15) was placed on top of each plate and then placed onto a shaker at 100 RPM for 20-min to dislodge the cells from the solid surface. The cell suspension was then funneled into a 15ml conical tube, pooling technical replicates (2 plates/replicate). The conical tube was then spun in the centrifuge at 5000 *RPM* for 3 minutes, to pellet cells. The supernatant was then poured off and decanted. Cells were then resuspended in 1 ml of PBS and placed in a 2 ml eppendorf tube and dyed with 1ul of already suspended “Life Technologies LiveDead Fixable Red 80 Assay” Dye from ThermoFisher (Catalog: L34971). The reaction was mixed by slight vortexing and allowed to incubate for 5 minutes, protected from the light. 1.1ml of 37% formaldehyde was then added to the reaction and left to fix the cells for 15 minutes, protected from the light. The fixed cells were then washed once with 1 mL of PBS with 1% bovine serum albumin, and finally resuspended in 1 mL of PBS with 1% bovine serum albumin. Fixed cells can be stored up to 3 days in the dark at 4**°**C before analysis on the flow cytometer. When ready for analysis, cells were filtered through a 5ml polystyrene round-bottom tube with 40uM cell strainer cap (falcon) and imaged on a BD LSR Fortessa (BD Biosciences). For each replicate, 20,000 cells were recorded and each experiment was performed at least three times. Cell death was imaged by using a 488 nm excitation laser and detected with a 685 LP 710/50 filter. Gates were set to exclude ungerminated conidia from the final analysis. Cells were categorized as having uptaken PI if fluorescence was detected above 10^3 in the PerCP-A channel. PI uptake percentages were plotted and analyzed in R. A more detailed protocol of the flow cytometry assay is detailed in **Supplementary File 1**.

### Confocal microscopy

Confocal microscopy was conducted in the Biology Imaging core facility at UC Berkeley. Specifically on the Carl Zeiss Inc LSM710 Confocal Microscope with differential Interference contrast (DIC) and GFP filters (400 nm excitation with 509 nm emission filter).

### RNA extraction

To extract total RNA, we carried out the same solid-surface interaction assay that was used in flow cytometry to phenotype vital dye uptake coupled to an RNA extraction protocol from Direct-zol RNA MiniPrep kit (Cat no. R2052) using the trizol/bead-beating method. In total, twelve samples, comprising four variables, pooling three technical replicates/samples were sequenced using Poly-A selection. The variables consisted of two timepoints: ten minutes and one hour and within each time point: bacterial exposure and negative control (MgCl_2_) exposure. After harvesting cells from the solid surface interaction assay, cells were placed in a twist cap tube with 0.1µM and 0.01µM glass beads and flash frozen with liquid nitrogen until ready to extract total RNA.

### RNA seq analysis

RNA was sequenced by Illumina (NovaSeq X Plus). The raw reads have been deposited to the SRA, BioProject accession number <u>PRJNA1209917</u>. Alignment, quantification and differential abundance analysis was done in KBase [89]. Raw reads were aligned to the OR74A (2489) *Neurospora crassa* genome using TopHat. HiSat2 was used to quantify reads into counts, and then FPKM values and TPM values. DEseq2 was used to calculate differential abundance of bacterial exposure to negative control. A standard P value cutoff of 0.05 [-log10(1.3)] and a fold change of >1 was selected. Descriptors were attached to the feature sets of differentially expressed genes, alongside the specific NCU#s (<u>Supplemental</u> <u>File 3</u>).

### Clustering and Enrichment analysis

To cluster genes based on their log fold change (log2FC) values, we employed the hclust function in R [90]. First, a distance matrix was computed using the Euclidean distance metric. The hierarchical clustering was performed using Ward’s method (Ward.D2), which minimizes the total within-cluster variance at each step. The clustering results were visualized as a dendrogram, with Euclidean distance represented on the y-axis. To identify distinct gene clusters, the dendrogram was cut at a height (h) of 7, resulting in the assignment of genes to 15 specific clusters. All computations and visualizations were performed using R. FunCat, GO and KEGG enrichment analysis was assigned to each cluster using Fungi-fun [[https://elbe.hki-jena.de/fungifun/] [74].

## Supplementary

1. **Supplementary** Figure 1: Exposure of *N. crassa* to PSTDC3000 and *E. coli* DH5a at a one-inch distance.
2. **Supplementary** Figure 2: Controls for solid-surface assay in Figure 2.
3. **Supplementary** Figure 3: Seven mutants of *N. crassa* that showed non-significant differences in growth.
4. **Supplementary File 1:** Flow Cytometry protocol for Bacterial-*N. crassa* interactions.
5. **Supplementary File 2:** CFU counting protocol of early time points of the *N. crassa* and PSTDC3000 interaction.
6. **Supplementary File 3:** Summary of each cluster in the enrichment analysis.
7. **Supplementary Table 1:** Data used for the creation of Figure 2.

a. Tab1: Figure 2C Surface area measurements.
b. Tab2: Figure 2E PSTDC3000 growth on differing *N.crassa* tissue types.
c. Tab3: Figure 2E PSTDC3000 hrcC mutant and *E. coli* DH5a growth on *N. crassa* hyphae.
8. **Supplementary Table 2:** Cell death measurements used for the creation of Figure 3.

a. Tab1: PI uptake percentages within the population of cells using flow cytometry (Figure 3C).
b. Tab2: Colony forming units (CFU/ml) measured (Figure 3D).
9. **Supplementary Table 3:** Intermediate files of RNAseq analysis

a. Tab1: Log_2_FC and P-values as calculated for all transcripts in N.crassa during this experiment.
b. Tab2: TPM as calculated for all transcripts in N.crassa during this experiment.
c. Tab3: FPKM as calculated for all transcripts in N.crassa during this experiment.
d. Tab4: 1,863 genes that were significantly differentially expressed (DE) fold change (Log_2_FC) ≥ 1, and (Log_2_FC) ≤ −1, false discovery rate (FDR) < 0.05.
e. Tab5: DE genes as clustered in Figure 4C.
f. Tab6: Genes of interest from enrichment analysis.
g. Tab7 - Tab21: FunCat, Kegg and GO analysis results for each individual cluster (as named on the tab).
10. **Supplementary Table 4:**

a. Tab1: Growth curves of all *N. crassa* mutants.
b. Tab2: Bacterial growth on *N. crassa* mutants.
c. Tab3: CFU measurements of *N. crassa* mutants.

## Data availability

SRA submission “Transcriptional sequencing of *Neurospora crassa* post *Pseudomonas syringae* DC3000 (PSTDC3000) inoculation, Dec 17’24” BioProject accession number <u>PRJNA1209917</u>. The 16S consensus sequences for this study can be found on GenBank using the accession number **SUB15006077.**

## Author Contributions

This project has been supported by the National Institute of Health Director’s Award (1DP2AT011967-01) awarded to Ksenia Krasileva, and UC Berkeley’s Chancellor Fellowship awarded to Frances Grace Stark. Research reported in this publication was supported in part by the National Institutes of Health S10 program under award number 1S10RR026866-01. The content is solely the responsibility of the authors and does not necessarily represent the official views of the National Institutes of Health. Frances Grace Stark conceptualized and executed the project with the help of Mari Torii-Karch and Sudyut Yuvaraj. Frances Grace Stark wrote the manuscript and synthesized all figures associated. Louise Glass and the Glass lab provided use of the full library of *N. crassa* OR74A mutants, and extensive feedback on this work. Pierre Glaudieux and Lucas Bonometti provided the environmental strains as well as extensive feedback.

## Acknowledgements

Thank you to the The RCNR Biological Imaging Facility at the University of California, Berkeley and Denise Schichnes for help in confocal microscopy techniques. Thank you Dr. Alexander Mela, Dr. Edyta Szewczyk and Dr. Monika Fischer for your extensive knowledge and help with Neurospora molecular genetics. We thank China Shaw for completing the admin associated with the collection of Neurospora’s bacteriome from the samples in France. Thank you to Dr. Edyta Szewczyk, Chandler Anne Sutherland, Jude Lange-Edwards, and Kyungyong Seong for extensive review and strengthening of this manuscript. The figures were created in BioRender (https://BioRender.com).

