## Supplementary Figure 1, Supplementary Figure 2, Supplementary Figure 3 for "Molecular Insights into Fungal Innate Immunity Using the *Neurospora crassa - Pseudomonas syringae* Model"

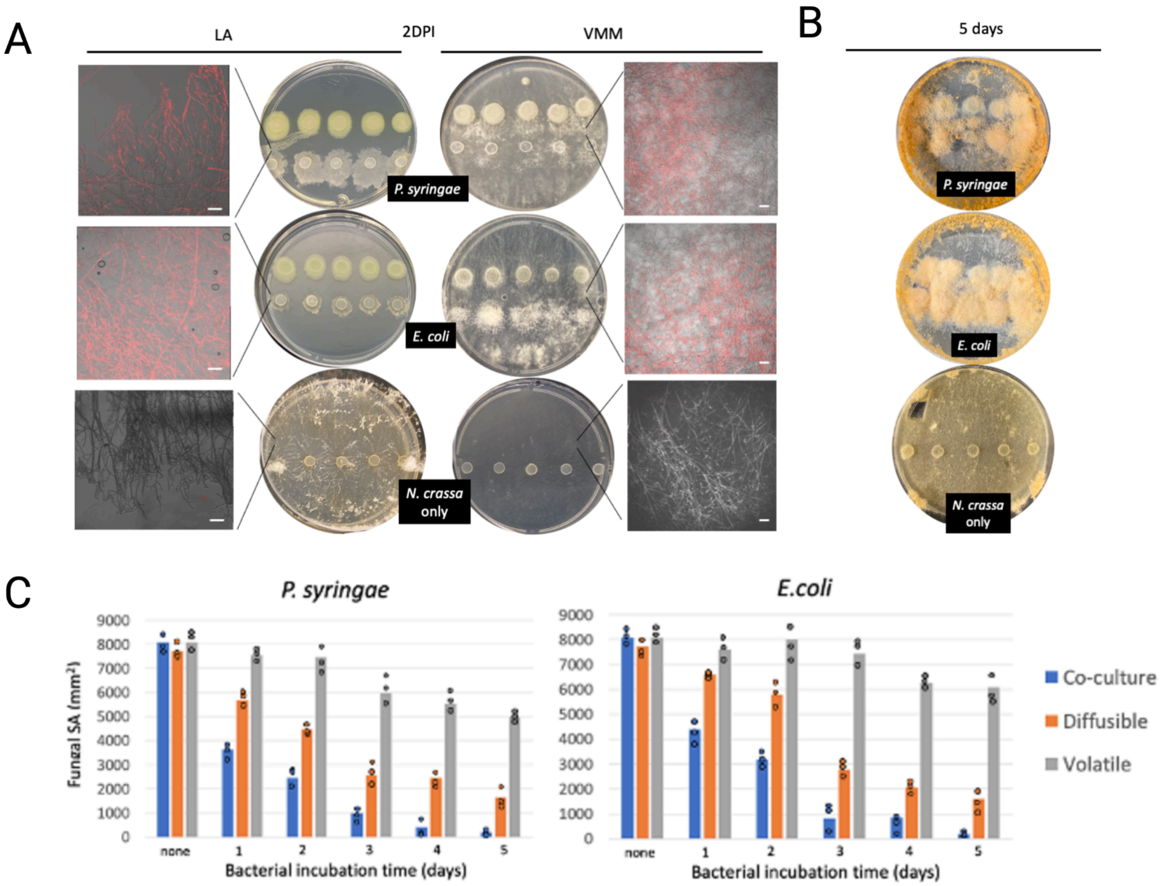


**Supplemental Figure 1: Exposure of *N. crassa* to PSTDC3000 and *E. coli* DH5a at a one-inch distance** A) Two days after bacterial exposure resulted in an alteration of fungal growth kinetics that pertained to the type of media- with nutrient rich Luria Agar (LA) showing defects and Nutrient poor Vogel's Minimal Medium (VMM) showing increased mycelium growth. Propidium Iodide (PI) uptake was observed in both VMM and LA 2 days post bacterial exposure. Scale bar= 100µm. B) *N. crassa* has increased conidiation on VMM seeded with bacteria five days after exposure. C) Co-culture, diffusible and volatile plate assays on LA revealed that growth defect is dependent on the dose of bacteria, and is mostly influenced by diffusible molecules.

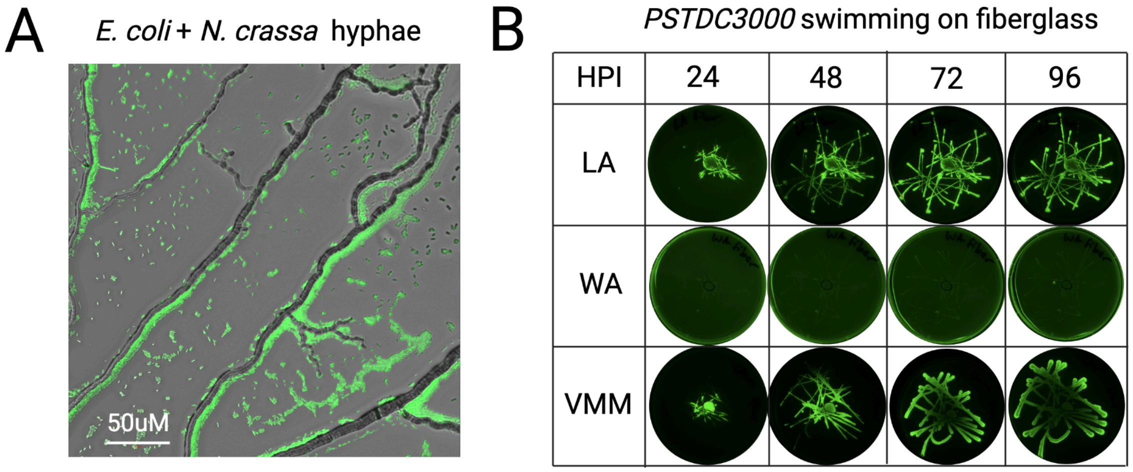


**Supplemental Figure 2: Controls for solid-surface assay.** A) *E. coli* DH5a also clusters on solid surfaces upon inoculum evaporation rather than the environment. B) PSTDC3000 was able to rapidly swim in all environments on fiberglass.

KO Mutants of *N.crassa* after exposure to *PSTDC3000*

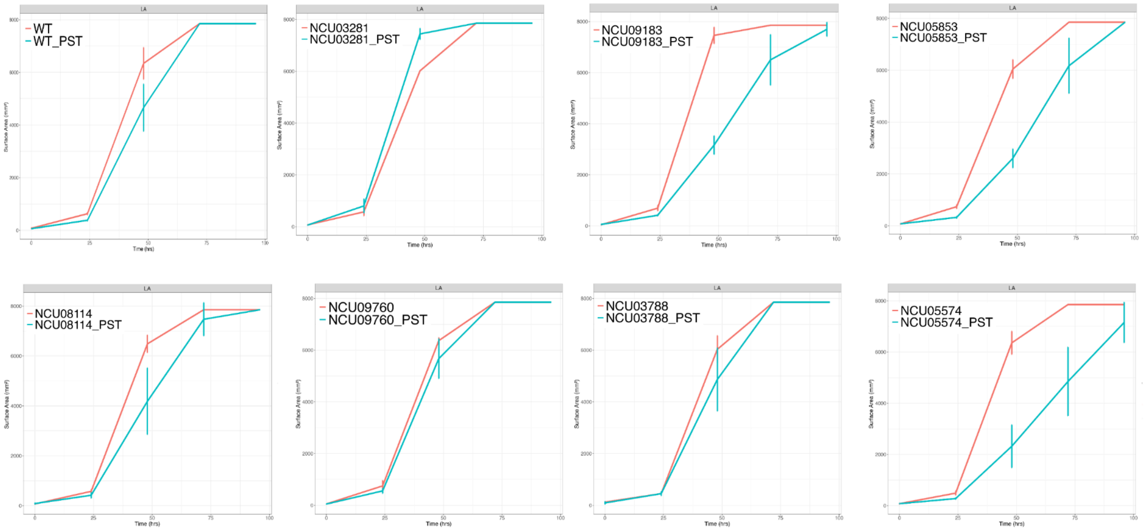


**Supplementary Figure 3: Seven mutants of *N. crassa* that showed non-significant differences.** Growth curves of mutants of N.crassa under PSTDC3000 pressure (n=6) that showed non-significant differences from Wild-Type using area under the curve (AUC) analysis coupled with paired T-tests.
