## Supplementary File 1 for "Molecular Insights into Fungal Innate Immunity Using the *Neurospora crassa - Pseudomonas syringae* Model"

### Flow Cytometry protocol for Bacterial- *N. crassa* interactions

Timeline example
- 5-7 days before experiment: Grow Neurospora strains
- Make sure you have a Flow Cytometry session booked on iLab (Can do after experiment)
- Day before the experiment: Grow overnight bacteria culture
- Day of Experiment
  - 9am: Inoculate N.crassa on water agar
  - 12pm: prep bacteria
  - 1:30 pm: Inoculate bacteria
  - 2:00 pm: Harvest
- Image on Flow Cytometer no more than 3 days post experiment

Variable examples
- Conidia (x3):
  - 2489 conidia
  - 2489 heat killed conidia
- Germlings (X3) per variable
  - 2489 WT Germlings + MgCl2 (24C)
  - 2489 WT germlings + *PSTDC3000* (24C)
- Bacteria:
  - *PSTDC3000*

Materials
- Cheesecloth (autoclaved)
- 15ml conical tubes, sterile
- 2 ml eppendorf tubes, sterile (preferred low-bind)
- Overnight bacteria in mid-log phase
- 5-7 day old conidia
- Water Agar with cellophane overlay
- PBS 1X
- 10mM MgCl2
- 15mM EDTA Ph6.4
- Life Technologies LiveDead Fixable Red 80 Assay dye from ThermoFisher (Catalog: L34971)
- 37% formaldehyde
- PBS 1X 1% BSA

Reconstituting the Life Technologies LiveDead Fixable Red Dye: (Skip if already done)

- 1.1 Bring one vial of the fluorescent reactive dye (Component A) and the vial of anhydrous DMSO (Component B) to room temperature before removing the caps.
- 1.2 Add 50 µL of DMSO to the vial of reactive dye. Mix well and visually confirm that all of the dye has dissolved.
- 1.3 Use the solution of reactive dye as soon as possible (see below), ideally within a couple days after reconstitution.
- Store reconstituted dye in the 20°C

#### Protocol for solid surface interaction assay

Overnight *PstDC3000* prep

- The day before the experiment take a test tube (20ml)
- Put 10mL of Lauria broth with a micropipetter into each test tube
- Put 10uL of Kanamycin and 10ul of Rifampicin into each test tube already containing LB
- Grab the *PstDC3000* plate from 4C that is no more than a month old
- With a micropipetter and a sterile pipette tip grab a small amount of bacteria from the plate.
- Eject the pipette tip containing bacteria into the LB + Kan + Rif test tube
- Place test tubes in the 30°C incubator (shaking) overnight.

*N. crassa* prep

- Grow Conidia for 5-7 days in VMM in a glass test-tube until a significant amount of macroconidiation has occurred.
- To harvest- 3ml of PBS directly into the tube and mix conidia until in solution.
- Put a sterile cheesecloth over a sterile 15mL conical tube.
- Filter the conidial suspension through the cheesecloth into the 15mL conical tube.
- Dilute conidia to 4.8 x 10^6 cfu/ml (OD of 0.3738).
    - Make enough in order to plate 500ul of conidia per plate (for germlings) and with at least 6ml left for conidial controls.
- Spread 500ul on the plates (cellophane plates- any environment) and let dry (w/lid open)
- Let germinate for 3.5- 4 hr at 30°C

Solid surface interaction assay

- look at the N. crassa plate under the microscope before starting the experiment to check if germination has occurred.
- If germinated, proceed with preparing the bacteria for inoculation

- Place 1ml of overnight bacterial suspension into the cuvette to get accurate reading of OD. Blank with liquid used to grow cells (LB).
- Multiply 500ul by the amount of plates getting bacterial treatment in order to calculate the mL of bacteria you need to make.
- Dilute the concentration of Bacteria to an OD600 of 0.5 = $5 \times 10^7$ cfu (colony forming units)/ml to your calculated volume.
- Pellet bacteria at 5000 RPM for 3 minutes, discard LB.
- Replace LB with the same amount of 10 mM $MgCl_2$ and resuspend bacteria by vortexing.
- use a cell spreader to inoculate 500ul of bacteria per plate over *Neurospora* seeded plates getting bacterial treatment (1:10 ratio) and just $MgCl_2$ for the negative controls.
- Let dry unto Neurospora for ~10min dry and incubate for however much time (30min average to get phenotype)

Harvesting the cells

- When ready to harvest, put 5ml 15mM EDTA (Ph = 6.4) on top of each plate using the macropipetter and glass pipettes. (doesn't have to be accurate, just over 5mL. More important to be fast).
- Put on a room temperature shaker for 10-20 min to dislodge the cells at 100 RPM
- Put a funnel on top of a sterile 15ml conical tube and dump harvested plates through the funnel. (For each variable have three 15ml conical tubes and pool two plates into one tube (total should be about 10ml)).
- Pellet the dislodged cells at 5000 RPM, for 3 minutes.
- Take off the supernatant (decant, but also try to get rid of any extra liquid), try not to disturb the pellet!

Fixing (*all centrifuge steps: 2 minutes at 10,000 x g)

- Resuspend pelleted cells in conical tube in 1ml of PBS
- Transfer 1ml to a 2ml eppendorf tube
- Add 1 µL of the reconstituted fluorescent reactive dye to 1 mL of the cell suspension and mix well by vortexing.
- Incubate at room temperature for 5min, protected from light.
- Add **110 µL** of 37% formaldehyde.
- Incubate at room temperature for 15 minutes, protect from the light
- Wash once with **1 mL** of PBS with 1% bovine serum albumin, and resuspend the cells in 1 mL of PBS with 1% bovine serum albumin.
- Store the tubes in 4*C in the dark to prevent photobleaching for up to three days before analysis

Flow Cytometry

- Grab the tubes from the 4*C
- Lightly mix them by pipetting, then filter 1 mL of the fixed cell mixture into a 5ml polystyrene round-bottom tube with 40uM cell strainer cap (falcon).
- Image on a BD LSR Fortessa (BD Biosciences). For each replicate: record 20,000 events.
- The dye is excited by a 488 nm laser and detected with a 685 LP 710/50 filter. Gates were set to exclude ungerminated conidia from the final analysis. Cells were deemed dead if fluorescence detected was above 10^3 in the PerCP-A channel. Cell death percentages were plotted and analyzed in R.
