## Supplementary File 2 for "Molecular Insights into Fungal Innate Immunity Using the *Neurospora crassa - Pseudomonas syringae* Model"

### Taking early time point CFU counts of *Neurospora crassa* and *PSTDC3000*

#### Timeline

1. 5-7 days before experiment: Grow Neurospora strains (in my experience fine to use up to ~15 day old conidia)
2. Day before the experiment: Grow overnight bacteria culture
3. Day of Experiment:
    - Inoculate N.crassa on water agar and grow for 3 hrs at 30°C + Prep bacteria to reach mid-growth curve in three hours
    - Inoculate bacteria 3hrs or when conidia have germinated
    - Harvest & plate after desired amount of time interacted

#### Protocol for solid surface interaction assay

Overnight *PSTDC3000* prep

- The day before the experiment take a test tube (20ml)
- Put 10mL of Lauria broth with a macropipetter into each test tube
- Put 10uL of Kanamycin and 10ul of Rifampicin into each test tube already containing LB
- Grab the *PstDC3000* plate from 4C that is no more than a month old
- With a micropipetter and a sterile pipette tip grab a small amount of bacteria from the plate.
- Eject the pipette tip containing bacteria into the LB + Kan + Rif test tube
- Place test tubes in the 30°C incubator (shaking) overnight.

*N. crassa* prep

- Grow Conidia for 5-7 days in VMM in a glass test-tube until a significant amount of macroconidiation has occured.
- To harvest put ~3ml of PBS directy into the tube and mix conidia until in solution.
- Put a sterile cheesecloth over a sterile 15mL conical tube.
- Filter the conidial suspension through the cheescloth into the 15mL conical tube.
- Dilute conidia to 4.8 x 10^6 cfu/ml (OD600 of 0.3738).
- Plate onto plated in 10ul drops. Can do up to 10 drops a plate
    - Suggest one variable/plate so all of the N + P on one plate, and all N + Mgcl2 (neg control on another plate)
- Let germinate for 3.5- 4 hr at 30°C
- Dilute overnight PST to OD 0.25- 0.3 OD600 in order to reach mid log phase by the time Neurospora is germinated

Solid surface interaction assay

- look at the N. crassa plate under the microscope before starting the experiment  to check if germination has occurred.
- If germinated, proceed with preparing the bacteria for inoculation
- Place 1 ml of overnight bacterial suspension into the cuvette to get accurate reading of OD. Blank with liquid used to grow cells (LB).
- Dilute the concentration of Bacteria to an OD600 of 0.5  ($5 \times 10^7$ cfu/ml)
- Pellet bacteria at 5000 RPM for 3 minutes, discard LB.
- Replace LB with the same amount of 10 mM  MgCl2 and resuspend bacteria by vortexing.
- Plate 10ul drop of PSTDC3000 directly on top of Neurospora 10ul drop. Add 10ul of MgCl2 for the negative controls and Pseudomonas on its own plate for Pseudomonas controls
- Let dry unto Neurospora for however much time (30 min average to get phenotype- suggest 2 hours)

Making CFUs

- When ready - cut out 1cm2 of each 10ul droplet and place in 1 ml of PBS in a 2 ml tube with 2 X 3.4mm STEEL beads
- Bead beat 1,500 RPM for 1 min
- Make a 1:10 dilution for all
- Add variables with Pseudomonas to 4 mg/ml AMP (Amphotericin b) to count Pseudomonas colonies
  - Suggest 20ul drops in three technical replicates
- Add variables with Neurospora to 2% sorbose + strep/gent to count Neurospora colonies
  - Place 100ul of 1:10 dilution onto sorbose and spread with a cell spreader
- Place both in the 24C incubator
- For negative controls
  - Place 10ul of starting conidia into 1ml of PBS - make a 1:10 dilution and plate

Counting CFUs

- Count Pseudomonas colonies the next day before they get too large
- Count Neurospora colonies 2 days post but much easier to see all colonies at 3 days
- Multiply by dilution factors after counting to get CFU/ml
