## Supplementary File 3 for "Molecular Insights into Fungal Innate Immunity Using the *Neurospora crassa - Pseudomonas syringae* Model"

### Summary of each cluster in the enrichment Analysis

**Clusters 2 and 15 consists of 10 min upregulated genes only.** Cluster 2 was enriched for genes associated with either transcription (rRNA synthesis/processing), translation and ribosomal biogenesis. Cluster 15 was a small cluster of only 3 genes, one as annotated as a HECT-type E3 ubiquitin transferase "thyroid hormone receptor interactor" (NCU09866T3) as well as other proteolytic activity such as dipeptidyl-peptidase-9 (*ddp-9*).

**Cluster 6 is shared upregulated genes:** Cluster 6 was enriched for C-compound and carbohydrate metabolism (polysaccharide metabolism) due to the inclusion of cell wall modifying enzymes, primary metabolism and TCA cycle proteins. Likewise, nitrogen metabolism was enriched. Enriched within transport facilities was heavy metal ion transport ($Cu^{+}$, $Fe^{3+}$, etc). GO enrichment also picked up "mating pheromone activity" & "positive regulation of mating-type specific transcription, DNA-templated" because of inclusion of the two isoforms of NCU01958, the *mat A-1* gene.

**Cluster 4 and 11 consists of 1 hr upregulated genes only:** Cluster 4 was enriched for protein folding stabilization/binding and included many heat shock proteins (Supplemental File 3), non-vascular ER transport alongside KEGG Protein processing in endoplasmic reticulum. There also seemed to be "mitotic chromosome condensation" and many proteins related to the condensin complex including kinesin-9 (NCU05180) which is a motor protein dictating microtubule binding and transport activity. There was also an upregulation of one transcript of the leashin Woronin body tether gene (NCU02793T3), which encodes the longest gene in *N. crassa* (~33KB).

**Cluster 14 consists of one gene** NCU10572T0, which encodes a predicted short chain oxidoreductase that is highly downregulated (Log2FC of -25.11) at 10 min and highly upregulated (Log2FC of 33.40) at 1 hr. The other transcript of NCU10572T1 is upregulated in cluster 6.

**Clusters 5,7 and 12 are downregulated clusters that capture clusters specific to 10min only:** Cluster five includes many metabolism and energy pathways. C-compound and carbohydrate metabolism (2.23E-07), Fatty acid metabolism (0.01) and lipid transport (0.04), biosynthesis of many amino acids: methionine, leucine and isoleucine, and two protective pathways under "Cell rescue defense and virulence" including a suggested glutathione conjugation reaction (0.002) and superoxide metabolism (0.002). Interestingly, the Cellular Component of GO enrichment was an “extracellular region” of which had many cellulose binding exoglucanases. Enriched in KEGG was secondary metabolism (1.27E-05). Cluster seven is a cluster consisting of 10min downregulated genes mating-type factor pheromone receptor activity (GO) as well as MAPK signaling pathway (KEGG) yeast were enriched because of the presence of pre-2 G-coupled Protein Receptor (GCPR) (NCU05758). FUNCAT also had meiotic recombination enriched by the presence of two isoforms of NCU01976. Cluster twelve consists of a short list of six genes incorporated processes of chromatin modification (FUNCAT, GO) and had transcription and transcription regulation, regulation of ARF protein signal transduction (NCU01465), and an NLR-like protein (NCU09760).

**Cluster 3,13 and 10 are downregulated clusters that are shared between the timepoints:** Cluster three consists mostly of shared genes that was highly enriched for nitrogen sulfate and selenium metabolism (1.65E-07), many fluxes of amino acid metabolism and oxidative stress response (0.02), Transport facilities, import, nonvascular import, ABC transporters, sulfate/sulfite, and anion transporters. The GO component had an integral component of membrane upregulated (7.37E-07) as indicated by the number of transporters within this cluster. Cluster five is a small cluster consisting of four shared downregulated genes. This included another transcript of the HECT-type E3 ubiquitin transferase “thyroid hormone receptor interactor” (NCU09866T2) and drug membrane antiporter activity (NCU04672). Cluster 10 consists of another transcript of the leashin Woronin body tether (NCU02793T2) that was highly downregulated at both time points. NCU02793T2 was in fact the most down-regulated gene in both independent timepoints.

**Clusters 1,8 and 9 are downregulated clusters capturing 1 hr only processes:** Cluster one is enriched with not as many hits as 10 min and continues to incorporate assimilation of ammonia, C-compound/carbohydrate metabolism, and transportation routes. NCU16992 pheromone *mfa-1*, transcription factor NCU07392 *adv-1* and another putative NLR-like protein (NCU08705T0) are found in this cluster. Cluster eight had no significant hits in FUNCAT and KEGG analysis. GO analysis revealed glycerophospholipid metabolism. Cluster 9 is highly downregulated at one hour and involves another transcript of the HECT-type E3 ubiquitin transferase “thyroid hormone receptor interactor” (NCU09866T1) alongside cellular component of vacuole, cytoplasmic vesicle, and MAPKKK signaling due to the presence of the cell wall integrity MAP-Kinase *mik-1* (NCU02234).
